# Bright Light Exposure Reduces Negative Affect and Modulates EEG Activity in Sleep-Deprived and Well-Rested Adolescents

**DOI:** 10.1101/2025.07.16.664355

**Authors:** Jana Kopřivová, Zuzana Kaňková, Přemysl Vlček, Marek Piorecký, Lenka Maierová, Zdeňka Bendová, Kateřina Skálová, Tereza Nekovářová

## Abstract

This study investigated whether a single morning session of bright light exposure modulates alertness, cognition, mood, and EEG activity in well-rested and partially sleep-deprived adolescents. Forty-seven subjects (15 – 21 years) were assigned to a well-rested (8 h sleep; 9 men, 15 women) or a sleep-deprived group (4 h sleep; 11 men, 12 women). All underwent 30 minutes of morning bright light exposure, with EEG, cognitive testing, and ratings of sleepiness and affect conducted pre- and post-intervention. Behavioral and electrophysiological changes were compared within and between groups. Associations between changes in EEG activity and behavioral outcomes were explored using correlation analyses. Bright light significantly reduced negative affect and improved Digit Span Forward task performance. No changes were observed in positive affect, subjective sleepiness, or Digit Span Backward scores. EEG analysis revealed decreased delta activity in the anterior cingulate cortex and increased beta activity in the right insula and fronto-parietal regions. Behavioral and EEG effects were similar across groups; however, only in the sleep-deprived group changes in beta activity significantly correlated with reduced negative affect. These results suggest that bright light may acutely enhance emotional state, cognitive performance, and cortical arousal in adolescents. The link between beta activity and affective improvement under sleep deprivation suggests a potential mechanism by which light supports emotional regulation.

## Introduction

Light plays a key role in shaping the behavior and physiology of organisms (Cajochen 2007; Vandewalle et al. 2009; LeGates et al. 2014). Environmental factors have long been recognized as central to psychophysical health even in archaic cultures (Novotná 2023). In addition to visual processing, light regulates several non-image-forming functions, including circadian rhythms, alertness, cognition, and mood (LeGates et al. 2014). These effects are mediated by intrinsically photosensitive retinal ganglion cells (ipRGCs), which transmit light information from the retina to the suprachiasmatic nuclei (SCN), the main circadian pacemaker, as well as to other brain regions involved in the regulation of sleep, cognition, and mood (LeGates et al. 2014). Although ipRGCs receive signals from rods and cones, they also contain their own photopigment, melanopsin, with a maximum sensitivity of about 480 nm (Hattar et al. 2002). This arrangement ensures that light in the blue spectrum, even at very low intensities, elicits a greater response in target structures than light absorbed by rods and cones (Gooley et al. 2010).

In recent years, research has increasingly focused on the non-image-forming effects of light, particularly its influence on alertness, cognitive function, and mood (e.g. Cajochen 2007; Vandewalle et al. 2009; Alkozei et al. 2021; Bjerrum et al. 2024). Daytime light exposure has been reported to enhance alertness and reduce sleepiness (Smolders & de Kort 2014; Kompier et al. 2021), with blue or blue-enriched light being more effective than longer wavelengths or light with lower color temperatures (Mills et al. 2007; Šmotek et al. 2019). Similarly bright white light showed a stronger alertness enhancing effect than dim light (Maierova et al. 2016; Phipps-Nelson et al. 2003; Rüger et al. 2006) but its effects are inconsistent in well-rested individuals when the control light level is at least 100 lux (Alwalidi and Hoffmann 2022). Cognitive benefits of bright light include improved attention, working memory, and reaction time (Okamoto and Nakagawa 2015), although effects vary with task complexity and time of day (Huiberts et al. 2015). EEG studies support these behavioral findings, showing that bright or blue light decreases low-frequency and increases high-frequency EEG power, indicating greater alertness (Šmotek et al. 2019; Zhang et al. 2024). Short-wavelength light also modulates cognitive processing, increasing P300 amplitude and reducing its latency (Okamoto and Kakagawa 2015; Šmotek et al. 2019).

Light exposure has also been consistently reported to impact mood. While a recent systematic review found that measures of affect were only influenced by narrow-band, long-wavelength light and reported mixed effects for white light, possibly due to high heterogeneity across studies (Bjerrum et al. 2024), several individual studies have reported that even a single light exposure can lead to measurable improvements in mood (Reeves et al. 2012; Leichtfried et al. 2015; Aan Het Rot et al. 2017). Blue-enriched light was found to be superior to standard office lighting in enhancing positive mood and reducing irritability (Viola et al. 2008). Mood improvement after a single exposure to bright light has been observed in individuals with seasonal affective disorder (Reeves et al. 2012), premenstrual mood disturbance (Aan Het Rot et al. 2017) and also in healthy subjects (Leichtfried et al. 2015). Moreover, neuroimaging studies have shown that blue light modulates brain networks involved in emotion regulation (Alkozei et al. 2021; Vandewalle et al. 2010).

Sleep-deprived individuals experience a stronger alerting effect in response to bright light (Alwalidi and Hoffmann 2022). Bright light has been shown to counteract cognitive decline and decrease sleep pressure in sleep-deprived subjects (Phipps-Nelson et al. 2003; Yokoi et al. 2003) and reduce afternoon decrease in alertness (Zhou et al. 2021). Sleep deprivation has consistently been shown to increase theta and delta EEG power (Münch et al. 2004), which can be suppressed by bright light (Zhang et al. 2024). Sleep deprivation also alters functional connectivity in the brain, specifically it reduces connectivity between the dorsolateral prefrontal cortex and the amygdala (Shao et al. 2014). Blue light may restore this connectivity (Alkozei et al. 2021), supporting its use to mitigate sleep deprivation effects and enhance mood regulation.

Adolescents face unique challenges related to light exposure due to both biological and environmental factors. Developmental changes in circadian physiology, such as delayed melatonin release (Crowley et al. 2007), coincide with increased evening exposure to light from electronic devices (Hale et al. 2018) and insufficient morning bright light exposure in indoor school environments (Figueiro et al. 2018), leading to reduced stimulation of non-image-forming visual pathways. In this context, daytime bright light exposure appears to be particularly beneficial for this specific age group, potentially improving mood regulation, alertness, and cognitive performance. While several studies have examined the effects of bright light on EEG activity, alertness, cognition, and mood in adults, there is limited evidence in adolescents. Long-term bright light exposure has been reported to improve alertness and cognition in this age group (Teicher et al. 2023) and has been suggested as a promising intervention to improve mood and sleep in adolescents (Holtmann et al. 2018). However, whether the immediate effects of bright light differ between sleep-deprived and well-rested adolescents remains an open question.

The purpose of this study was to investigate how a single morning session of bright light exposure modulates alertness, cognitive performance, mood, and EEG patterns in well-rested and partially sleep-deprived healthy adolescents. We included both well-rested and sleep-deprived groups to examine whether the non-image-forming effects of light differ depending on sleep status, and whether bright light may mitigate the consequences of sleep loss.

## Materials and Methods

### Design

This study examined the effects of a 30-minute bright light exposure in adolescents pseudo-randomly assigned either to the non-deprived (ND, 8 ± 1 hours of sleep) or the sleep-deprived (SD) group who slept 4 hours with a delayed bedtime. The following morning, the participants were tested at the National Institute of Mental Health with cognitive tests, self-reported scales, and underwent EEG examinations. All measures were taken twice, before and immediately after light exposure that occurred between 10:00 and 12:00 noon. This time window was selected to reflect a typical mid-morning school period when adolescents are exposed to relatively low indoor light levels. The interval between the first and second EEG measurement was approximately 1 hour (64 ± 10 min), the EEG cap was not removed between the two measurements. To minimize the potential confounding effects of seasonal variations in daylight, data collection was restricted to the standard time period (February–March 2022, October–December 2022, and March 2023). The study was approved by the local Ethics Committee of the National Institute of Mental Health. All participants provided written informed consent; for those under 18, consent was also obtained from their legal guardians. The study was conducted in accordance with the Declaration of Helsinki.

### Participants

Fifty-five healthy adolescents participated aged 15 – 21 years were enrolled in the study. Exclusion criteria included any personal or first-degree family history of psychiatric disorders, a history of head injury resulting in loss of consciousness, any active or untreated medical condition (including neurological or cardiovascular diseases), current or recent substance use problems, neurological disease or MRI-detected brain pathology, any diagnosed chronic sleep disorder or acute sleep problems during the week preceding the study, regular use of medication affecting sleep or alertness, and daily caffeine consumption exceeding 300 mg. Eligibility was determined based on participants’ self-report responses to the inclusion and exclusion criteria, and for those under 18 years, confirmed by their legal guardians. Eight subjects were excluded due to missing or poor-quality EEG recordings at either the first or second measurement. Of the remaining 47 subjects, 24 were part of the ND group and 23 of the SD group. The final sample consisted of 20 males and 27 females aged 15 to 21 years (median: 16 years, interquartile range: 16-19 years). The SD and ND groups were matched for age (p = .596) and sex (SD group: 11 men, ND group 9 men, p = .561). Although participants were not matched for chronotype, screening data from a larger sample (n = 99) from which the study participants were recruited indicated that most individuals had an intermediate chronotype (76/99), and none exhibited an extreme chronotype.

### Behavioral and cognitive measures

Subjectively experienced emotions were assessed using the Positive and Negative Affect Scale, PANAS (Watson et al. 1988). The PANAS consists of two 10-item scales measuring positive and negative emotions. Each item is scored from 1 to 5. The Stanford Sleepiness Scale, SSS (Hoddes et al. 1972), a single-item self-report questionnaire with a seven-point scale, was administered to assess sleepiness. Cognitive function was assessed using the Digit Span test (Wechsler 2013) which includes forward (DSF) and backward (DSB) number repetition. The forward task primarily measures short-term memory and attention, while the backward task assesses working memory and executive functions such as cognitive control and inhibition (Wechsler 2013).

### Electroencephalography

Resting-state EEG was recorded using a Brainscope amplifier (M&I, Czech Republic) from 21 electrodes placed according to the international 10–20 system (sampling rate: 1 kHz, band-pass: 0.1–200 Hz). Each recording lasted 20 minutes (10 min eyes open, 10 min eyes closed), but only eyes-open data were analyzed to reduce vigilance-related variability. Prior to analysis, all electroencephalographic (EEG) recordings underwent rigorous pre-processing by an experienced rater using Brain Vision Analyzer software (Brain Products GmbH, Germany). The data were band-pass filtered between 1 and 60 Hz, with a notch filter applied at 50 Hz to eliminate power line interference. Ocular artifacts were corrected using Independent Component Analysis (ICA), ensuring the removal of eye movement–related contamination. Additional artifacts, including muscle-related noise and other non-neural interferences, were identified and removed through a meticulous visual inspection by an experienced rater. Only artifact-free, two-second epochs were retained for subsequent quantitative analysis, ensuring the integrity of the dataset for further processing. Data were analyzed using the eLORETA (exact low resolution brain electromagnetic tomography) method in the LORETA-Key software (http://www.uzh.ch/keyinst/loreta.htm). eLORETA is a linear inverse solution for estimating the intracranial distribution of current density in three-dimensional space by reconstructing the intracerebral electrical sources underlying scalp-recorded activity using a three-shell spherical head model co-registered with Talairach coordinates (Pascual-Marqui 2002; Pascual-Marqui 2007; Pascual-Marqui et al., 2011). eLORETA estimates the current density distribution in 6,239 voxels with a spatial resolution of 5 × 5 × 5 mm and is restricted to cortical grey matter and hippocampi. In this study, current density values were computed in six frequency bands: delta (2 - 4 Hz), theta (4.5 - 8 Hz), alpha (8.5 - 12 Hz), beta 1 (12.5 - 21 Hz), beta 2 (21.5 - 30 Hz), gamma (30.5 - 45 Hz).

### Light pavilion

The indoor light pavilion is a cubial space with a side length of 2.5 m and seating for up to six people. The entire ceiling and half of one side wall are fitted with LED light sources with a diffuser so that the whole area acts as a flat light source with balanced brightness. The large, sky-like light surface provides glare-free illumination. The system emits balanced-spectrum light (4500 K, Ra > 80) with up to 11,500 lx corneal photopic illuminance, corresponding to >11,000 lx melanopic daylight equivalent (mEDI, International Commission on Illumination 2018). In this study, 8000 lx (7500 lx mEDI) was used. The target mEDI level was derived from the commonly recommended photopic illuminance for bright-light therapy (10000 lx) and the spectral properties of the fluorescent light device SAD 6000 (Waldmann Medizintechnik, Germany), one of the fluorescent light boxes used in clinical practice during the early era of light therapy. The corresponding mEDI value was calculated according to the CIE S 026/E:2018 standard (International Commission on Illumination, 2018) using the CIE Toolbox S 02 (https://bit.ly/2T9QLTL). The lighting technology used in this study (Spectrasol, Czech Patent No. 308363) features a full spectral band with enhanced power in the 460–520 nm range, ensuring strong activation of ipRGCs and non-visual photoreception (Fig. 1). Owing to its enhanced spectral power within the melanopically sensitive range, this light source reaches 7500 lx mEDI already at 8000 lx photopic illuminance.

**Figure 1.**
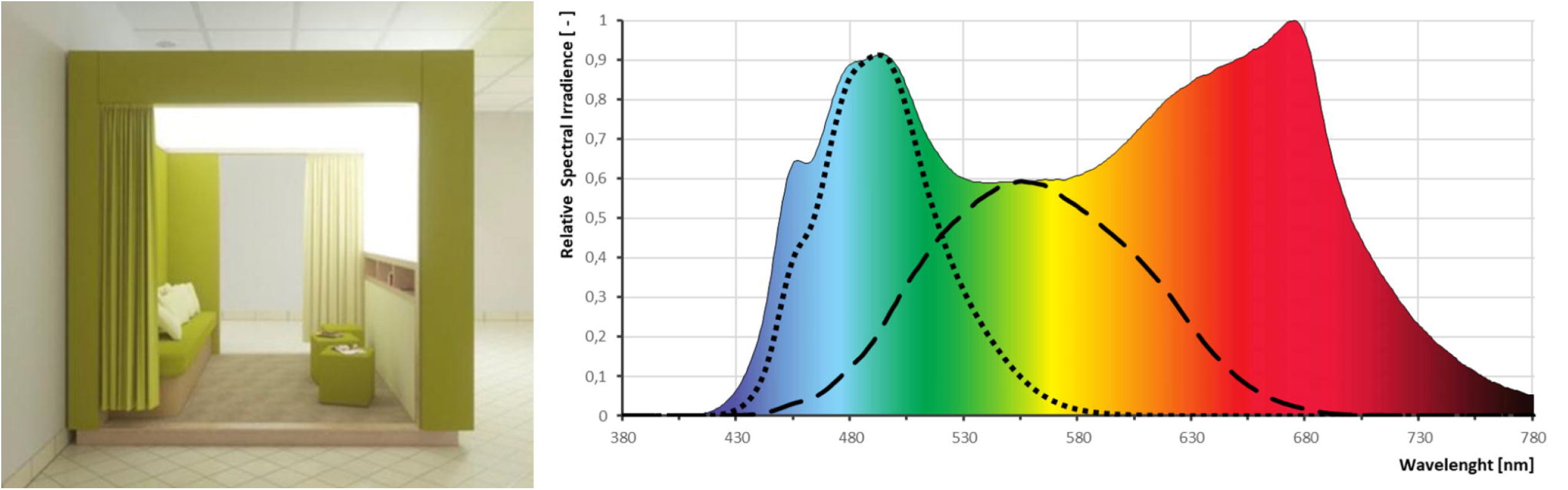
The indoor light (left) pavilion and characteristics of the light (right). The colored area represents the spectral power distribution of the light source. Dashed and dotted lines indicate the spectral sensitivity functions for photopic illuminance (black dashed line) and melanopic illuminance (black dotted line), respectively.

**Figure 2.**
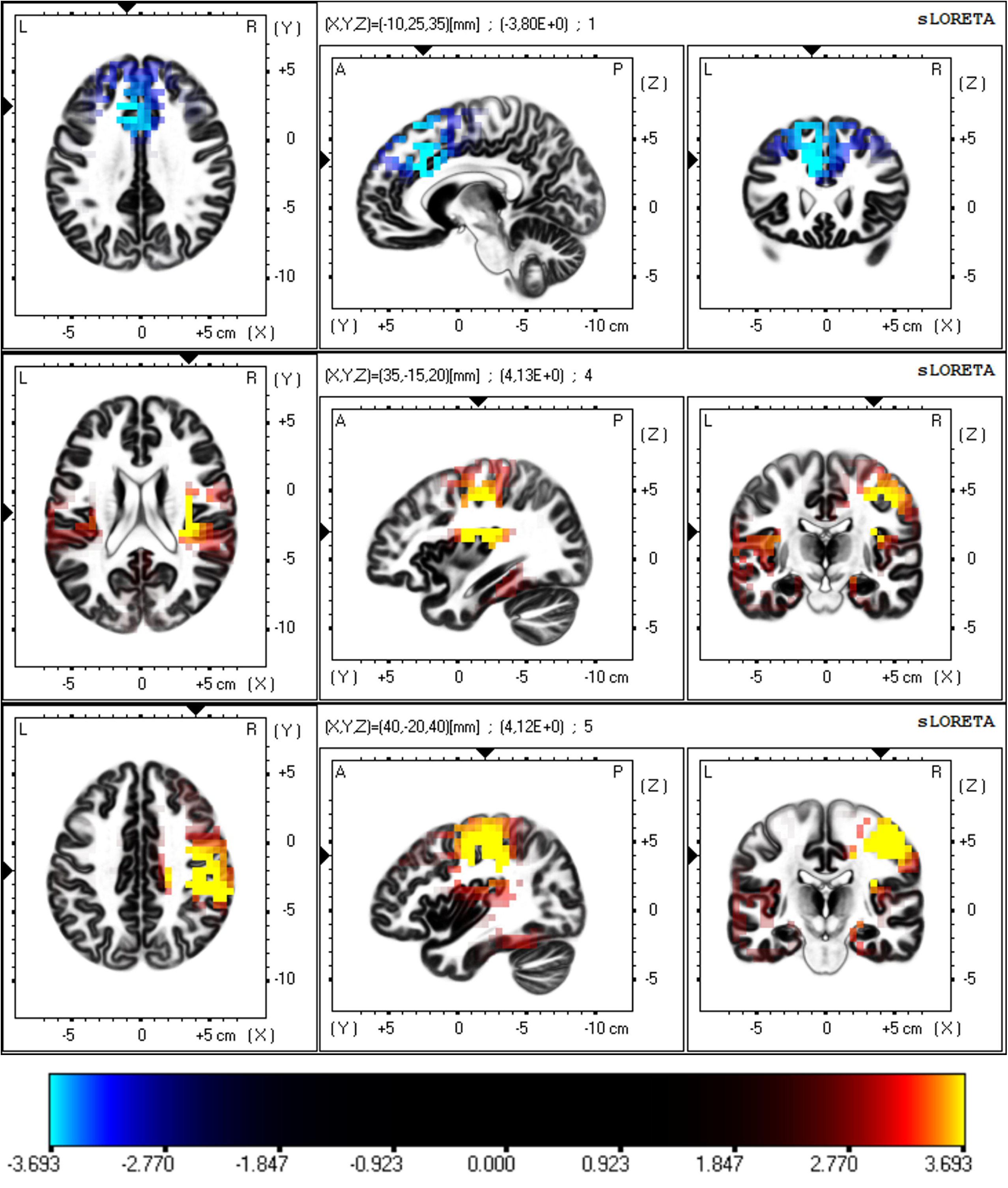
Difference in EEG activity assessed by eLORETA before and after light exposure. The voxels with lower or higher EEG activity in the given frequency bands after light exposure are marked in cyan and yellow, respectively (p < 0.05, corrected). Each figure is sliced to its own maximum value of the permutation-derived threshold (3.919). The scale exponent was set to 5 to enhance visibility of significant and near-threshold voxels while maintaining figure clarity. Abbreviations: eLORETA – exact low-resolution brain electromagnetic tomography.

### Statistical analysis

#### Behavioral data

Since most of the variables did not meet the assumption of normality, primary analyses were conducted using non-parametric tests (Wilcoxon signed-rank and Mann–Whitney U tests). Values that were more than 3 interquartile ranges (IQRs) away from the median were considered extreme outliers and excluded from the analysis. Outlier detection was performed across all participants for each variable separately. A total of four outliers were identified across all behavioral variables, two in the SSS scale scores (1 subject from the SD group) and two in the DSB test scores (2 different subjects from the ND group). To assess the robustness of the findings and to examine potential interaction effects, we additionally conducted a two-way robust ANOVA using Yuen’s method with 20% trimmed means. This approach accounts for violations of normality and the presence of outliers, which were retained in the analysis.

#### EEG data

Since the EEG data analyses were paired, it was not necessary to perform normalization of the EEG signal, which is usually required in intergroup comparisons due to interindividual differences in EEG amplitude. Absolute eLORETA current density values for each voxel were compared using the Wilcoxon signed rank test in the LORETA-KEY software. Non-parametric statistical mapping with 5,000 permutations was used to determine statistical significance (Nichols and Holmes 2002). All bands were treated simultaneously to ensure that the family-wise type I error did not exceed the nominal level p = .05 after correction for multiple comparisons across all voxels and all frequencies.

#### Correlation between EEG and behavioral data

To test possible relationships between EEG changes and changes in mood and cognitive performance, the mean difference in current density (post-intervention minus baseline) in cortical voxels that showed significant differences before and after the intervention was correlated with the difference scores (post-intervention score minus baseline score) on the behavioral scales that showed distinct values after the intervention compared to baseline. As our data were not normally distributed, the Spearman correlation coefficient was calculated. Values more than 3 IQRs from the median were considered extreme outliers. In total, four outlying values were found in the EEG current density differential score (two values in the delta band — one from the SD and one from the ND group — and one value in the beta 1 and one in the beta 2 frequency band, both from the same subject belonging to the SD group) and were removed before correlation analysis was performed.

## Results

### Behavioral and cognitive measures

The intervention significantly reduced negative affect in the whole cohort, with no change in positive affect or sleepiness. Similar results were found for both subgroups: negative affect decreased in the ND as well as in the SD group; positive affect and sleepiness remained unchanged. DSF performance improved in the whole group and in the ND subgroup; significance was not reached in the SD subgroup. No significant change was observed in DSB performance across groups (Table 1). At baseline, the SD group showed higher levels of sleepiness and worse performance on the DSF test. Post-intervention, the difference in sleepiness was no longer observed, but the ND group still outperformed on the DSF task. However, differential scores (post minus baseline) did not differ between groups (see Table 2). To further examine whether bright light exposure could alleviate the effects of sleep deprivation, we compared post-intervention outcomes in the SD group to baseline values of the ND group using the Mann–Whitney U test. The SD group remained significantly sleepier (SSS: U = 177, p = .041) than the ND group at baseline, whereas no significant differences were found for affect (PANAS-P, PANAS-N) or cognitive performance (DSF, DSB; all p > .320).

**Table 1.** Pre–post changes in affect, sleepiness, and cognitive performance in the total sample and subgroups. The table reports medians and interquartile ranges (IQR) at baseline and after the light intervention, along with Wilcoxon signed-rank test statistics (Z), p-values, and effect sizes (r) for within-group comparisons. Abbreviations: ND – non-deprived group; SD – sleep-deprived group; PANAS-P – Positive Affect subscale of the Positive and Negative Affect Schedule; PANAS-N – Negative Affect subscale of the Positive and Negative Affect Schedule; SSS – Stanford Sleepiness Scale; DSF – Digit Span Forward; DSB – Digit Span Backward; r – effect size (rank-biserial correlation).

| Measure | Group | Baseline (Median, IQR) | Post-intervention (Median, IQR) | Pre-post difference (Wilcoxon signed-rank test) |  |  |
| --- | --- | --- | --- | --- | --- | --- |
|  |  |  |  | Z | p | r |
| PANAS-P | All | 29 (24 - 33) | 27 (22 - 32) | 1.56 | .120 | 0.264 |
|  | ND | 29.5 (24.25 - 32.75) | 27 (22.25 - 30.75) | 1.13 | .264 | 0.268 |
|  | SD | 29 (21 - 34) | 28 (20 - 33) | 1.00 | .322 | 0.239 |
| PANAS-N | All | 16 (12 - 20) | 13 (10 - 15) | 4.28 | < .001 | 0.768 |
|  | ND | 14.5 (12 - 17) | 11.5 (10 - 14) | 3.38 | < .001 | 0.844 |
|  | SD | 18 (13 - 28) | 13 (11 - 20) | 2.83 | .005 | 0.706 |
| SSS | All | 3 (2 - 3) | 3 (2 - 3) | 0.38 | .690 | 0.080 |
|  | ND | 2 (2 - 3) | 2 (1 - 3) | 0.60 | .549 | -0.181 |
|  | SD | 3 (2.75 - 4) | 3 (2 - 3) | 1.16 | .217 | 0.331 |
| DSF | All | 10 (9 - 12) | 11 (9 - 13) | 3.26 | < .001 | -0.640 |
|  | ND | 11 (9 - 13) | 12.5 (10.25 - 14) | 2.83 | .004 | 0.760 |
|  | SD | 9 (8 - 12) | 10 (9 - 12) | 1.78 | .070 | -0.500 |
| DSB | All | 7 (6 - 8) | 7 (6 - 8) | 1.58 | .107 | -0.320 |
|  | ND | 7 (6 - 9) | 8 (7 - 8) | 1.28 | .199 | -0.375 |
|  | SD | 7 (6 - 8) | 7 (5 - 8) | 0.95 | .344 | -0.261 |

**Table 2.** Behavioral characteristics of the non-deprived (ND) and sleep-deprived (SD) subgroups. The table shows medians and interquartile ranges (IQR) for each group at baseline, after the intervention, and for differential scores (post–pre). Between-group comparisons were conducted using the Mann–Whitney U test; test statistics (U), p-values, and effect sizes (r) are reported. Abbreviations: ND – non-deprived group; SD – sleep-deprived group; PANAS-P – Positive Affect subscale of the Positive and Negative Affect Schedule; PANAS-N – Negative Affect subscale of the Positive and Negative Affect Schedule; SSS – Stanford Sleepiness Scale; DSF – Digit Span Forward; DSB – Digit Span Backward; r – effect size (rank-biserial correlation).

|  | ND group (n = 24) | SD group (n = 23) | SD vs. ND difference (Mann-Whitney U test) |  |  |
| --- | --- | --- | --- | --- | --- |
|  |  |  | U | p | r |
| Baseline measurement |  |  |  |  |  |
| PANAS-P | 29.5 (24.25 - 32.75) | 29 (21 - 34) | 264.50 | .814 | -0.042 |
| PANAS-N | 14.5 (12 - 17) | 18 (13 - 28) | 347.50 | .130 | 0.259 |
| SSS | 2 (2 - 3) | 3 (2.75 - 4) | 382.00 | .007 | 0.447 |
| DSF | 11 (9 - 13) | 9 (8 - 12) | 172.50 | .035 | -0.357 |
| DSB | 7 (6 - 9) | 7 (6 - 8) | 229.00 | .434 | -0.134 |
| Post-intervention measurement |  |  |  |  |  |
| PANAS-P | 27 (22.25 - 30.75) | 28 (20 - 33) | 269.50 | .898 | -0.024 |
| PANAS-N | 11.5 (10 - 14) | 13 (11 - 20) | 360.50 | .070 | 0.306 |
| SSS | 2 (1 - 3) | 3 (2 - 3) | 322.50 | .186 | 0.222 |
| DSF | 12.5 (10.25 - 14) | 10 (9 - 12) | 163.50 | .016 | -0.408 |
| DSB | 8 (7 - 8) | 7 (5 - 8) | 216.50 | .287 | -0.181 |
| Differential score (post-intervention vs. baseline measurement) |  |  |  |  |  |
| PANAS-P | -0.5 (-5.5 - 1.75) | -1 (-8 - 3) | 269.00 | .890 | -0.025 |
| PANAS-N | -2 (-4 - -0.25) | -2 (-7 - 0) | 290.00 | .773 | 0.051 |
| SSS | 0 (-0.75 - 1) | -1 (-1 - 0) | 204.00 | .111 | -0.261 |
| DSF | 1 (0 - 2) | 1 (0 - 2) | 250.50 | .587 | -0.092 |
| DSB | 0 (-1 - 1.25) | 0 (-1 - 1) | 268.50 | .879 | -0.027 |

**Table 3.** Differences in EEG activity before and immediately after a 30-minute bright light exposure. The table lists the areas where significant differences (decrease or increase in EEG activity) were found before and after the intervention. The number of significant voxels for each region and the frequencies and mean permutation-derived test statistics are indicated.

|  | Delta (2 - 4 Hz) |  | Beta 1 (12.5 - 21 Hz) |  | Beta 2 (21.5 - 30 Hz) |  |
| --- | --- | --- | --- | --- | --- | --- |
|  | mean test stat. | voxels | mean test stat. | voxels | mean test stat. | voxels |
| **Brain structure and Brodmann area (BA)** |  |  |  |  |  |  |
| Anterior Cingulate Gyrus (BA 32) | -3.730 | 8 |  |  |  |  |
| Anterior Cingulate Gyrus (BA 24, 32) | -3.721 | 3 |  |  |  |  |
| Medial Frontal Gyrus (BA 6, 8, 9) | -3.711 | 3 |  |  |  |  |
| Insula (BA 13) |  |  | 3.894 | 14 | 3.771 | 3 |
| Inferior Parietal Lobule (BA 40) |  |  | 3.817 | 3 | 3.843 | 26 |
| Precentral Gyrus (BA 4, 6) |  |  | 3.804 | 18 | 3.814 | 46 |
| Postcentral Gyrus (BA 1, 2, 3, 40) |  |  | 3.798 | 16 | 3.855 | 65 |
| Middle Frontal Gyrus (BA 6) |  |  | 3.743 | 3 | 3.733 | 5 |

Trimmed ANOVA analyses partially supported the findings from non-parametric tests. For positive affect, no significant main effects of group (p = .807), time (p = .277), or group × time interaction (p = .811) were observed. For negative affect, a significant main effect of group (Q = 4.80, p = .034) and time (Q = 9.32, p = .004) was found, without a significant interaction (p = .752). Post-hoc comparisons revealed that the SD group had overall higher scores than the ND group (ψ[ = 4.50, p = .034, 95% CI [0.36, 8.65]) and that there was a significant reduction over time across groups (ψ[ = 6.28, p = .004, 95% CI [2.14, 10.40]). For subjective sleepiness, a significant group effect was found (Q = 7.64, p = .008), indicating higher levels in the SD group regardless of time (time: p = .613; interaction: p = .336). This difference was confirmed by post-hoc analysis (ψ[ = 1.49, p = .008, 95% CI [0.41, 2.58], see Supplementary Figure 1). DSF scores showed significant main effects of both group (Q = 12.53, p = .001) and time (Q = 4.42, p = .040), indicating better performance in the ND group and improvement following the intervention in the whole sample. Post-hoc tests confirmed the group difference (ψ[ = −3.65, p < .001, 95% CI [−5.72, −1.59]) and the pre–post improvement (ψ[ = −2.17, p = .040, 95% CI [−4.24, −0.10]). No significant group × time interaction was found (p = .498). For DSB, no significant effects were observed for group (p = .177), time (p = .411), or interaction (p = .802).

### Electrophysiological data

In the whole cohort, light exposure led to decreased EEG activity in the delta band and increased activity in the beta 1 and beta 2 bands. Delta reductions were localized mainly in the left anterior cingulate cortex (BA 32), while beta increases were found in the right insula (BA 13) and fronto-parietal regions including the inferior parietal lobule and sensorimotor cortex. For beta 1 the largest differences were observed in the right insula (BA 13), for beta 2 the largest differences were found in the inferior parietal lobule (BA 40; p < .05, corrected; Fig. 3). Detailed statistics are provided in Table 2.

Importantly, the magnitude of EEG changes did not differ significantly between the SD and ND groups: for delta, U = 291.00, p = .398, r = 0.150, SE = 0.172; for beta 1, U = 232.00, p = .492, r = 0.121, SE = −0.171; and for beta 2, U = 255.00, p = .853, r = −0.034, SE = 0.171.

A separate analysis revealed no significant differences in eLORETA current density between the SD and ND groups before and after the intervention. However, voxel-wise trends observed in the whole group (increased beta, decreased delta) were also present in both subgroups, with post-intervention values shifting in the same direction as in the overall cohort.

### Correlation analysis

In the SD group, a greater increase in beta 1 activity in the right insula, frontal, and parietal sources was significantly associated with a greater reduction in negative affect (Spearman’s ρ = −0.434, p = .043). In contrast, the ND group showed no such association (ρ = −0.009, p = .966), and in the whole cohort, the correlation approached significance (ρ = −0.268, p = .072). Beta 2 showed a similar negative trend in the SD group (ρ = −0.340, p = .121). No significant associations were found between EEG changes and DSF performance in any group. Similarly, changes in delta activity did not show significant correlations with changes in negative affect (ρ = −0.088, p = .566 in the whole group; ρ = −0.126, p = .576 in the SD group; ρ = −0.040, p = .857 in the ND group) or with changes in DSF performance (ρ = 0.094, p = .539 in the whole group; ρ = 0.046, p = .839 in the SD group; ρ = 0.140, p = .525 in the ND group). A formal comparison of independent correlations for the beta 1 – PANAS-N relationship between the SD and ND groups using Fisher’s z revealed that the difference between the two groups was not statistically significant (z = 1.50, p = .134).

## Discussion

A thirty-minute stay in the light pavilion with exposure to bright light reduced negative affect, improved DSF performance, suppressed slow and enhanced fast frequencies in the EEG. Sleep-deprived subjects showed greater sleepiness and worse performance on the DSF task at baseline compared to well-rested subjects. Overall, the data suggest that the light intervention had similar behavioral and electrophysiological effects in both SD and ND groups. Individual correlations observed only in the SD group indicate a potential mechanism through which light supports emotional regulation.

Our study revealed a decrease in delta power in the dorsal anterior cingulate cortex (dACC) following exposure to bright light. The dACC plays a critical role in cognitive control and attentional regulation (Bush et al., 2000). However, although increased EEG delta activity has previously been associated with reduced vigilance, theta activity showed a more specific relationship to individual attentional lapses (Makeig and Jung, 1996). Our finding of decreased delta power in the dACC following bright light exposure is consistent with recent studies employing daytime exposure to bright light and repeated resting-state EEG measurements. Lok et al. (2018) and Luo et al. (2021) both reported significant reductions in delta activity when participants were exposed to 1200 lx versus 200 lx polychromatic white light for 5 hours in the afternoon. Notably, both studies found that theta activity remained unaffected by light condition, mirroring our findings and suggesting that daytime bright light selectively modulates delta activity rather than broadly suppressing all low-frequency EEG rhythms. These findings contrast with an earlier study by Badia et al. (1991), which found no significant EEG changes during 90-minute daytime exposure to bright light (5000-10000 lx vs. 50 lx) in the afternoon. However, it is important to note that all three of these studies assessed EEG during light exposure, whereas our study focused on comparing EEG before and after light exposure. The observed reduction in delta power in the dACC following bright-light exposure should therefore be interpreted with caution and requires replication in independent samples using comparable daytime resting-state protocols. Our results show that a 30-minute exposure to bright light increased beta-1 and beta-2 activity which is consistent with previous daytime light exposure studies (Lok et al. 2018; Luo et al. 2021). The significant sources were localized in the right insula and fronto-parietal cortical regions, including the inferior parietal lobule, and pre-/postcentral gyrus, which corresponds with previous studies showing light-induced activation in the insula and regions near the intraparietal sulcus (Chen et al. 2021). The insula, particularly its right anterior region, is involved in salience detection, interoception, and integration of autonomic and cognitive functions (Molnar-Szakacs and Uddin 2022). Similarly, the parietal cortex, specifically Brodmann area 40 including the inferior parietal lobule, plays a critical role in multisensory integration and cognitive processing (Corbetta and Schulman 2002). Thus, increased beta activity in these areas may reflect enhanced cognitive regulation and behavioral readiness. Although the insula does not receive direct projections from the ipRGCs, it connects with the lateral habenula (Vadovičová 2014), a structure involved in negative affect regulation (Matsumoto and Hikosaka 2007) and directly innervated by these retinal cells (LeGates et al. 2014).

Indeed, our study showed that a thirty-minute stay in the light pavilion with exposure to bright light reduced negative affect in both the SD and ND subgroups, while positive affect remained unchanged. This aligns with previous research demonstrating that even a single session of blue or bright light can improve mood, particularly by reducing negative affect. For instance, Alkozei et al. (2021) reported that blue light increased amygdala–prefrontal connectivity, correlating with mood improvement. Similar effects were observed in women with premenstrual dysphoria (Aan Het Rot et al. 2017) and during acute tryptophan depletion, where bright light prevented mood deterioration (Aan Het Rot et al. 2008).

The increase in beta activity after light exposure correlated with a reduction in negative affect only in sleep-deprived individuals. Although the between-group difference in correlation coefficients did not reach statistical significance, this pattern may indicate that the neurophysiological processes engaged by light exposure are more tightly associated with affective changes under sleep deprivation. Sleep and emotional brain function are closely linked (Goldstein and Walker 2014). Self-reported sleep duration has been negatively related to subjective psychological distress and prefrontal-amygdala connectivity (Killgore 2013). Sleep deprivation has been shown to affect the communication network between the prefrontal cortex and limbic structures and to adversely modulate the brain’s response to negative aversive stimuli (Yoo et al. 2007).

Interestingly, blue light has been found to strengthen functional connectivity between the dorsolateral prefrontal cortex and the amygdala, and the increase in connectivity between these two regions was associated with a reduction in negative affect (Alkozei et al. 2021). Although our study did not specifically assess functional connectivity, the increased beta activity observed after the intervention may indicate the restoration of certain aspects of emotional regulation in response to bright light.

This study has several limitations. First, participants’ recent light exposure history was not controlled, which may have contributed to interindividual variability in responses to bright light. However, as all participants were high-school students with regular morning classes starting at 8:00 a.m., their morning light exposure routines were likely similar. Second, the absence of a control light condition limits causal interpretation, as reductions in negative affect may reflect decreasing anxiety over time or the social context of the pavilion. Although interaction occurred before measurements, the social environment during light exposure may have influenced results. The study design also does not fully exclude practice effects on cognitive performance. However, the improvement was limited to DSF, which is relatively resistant to short-term practice, while no gains were observed in the more practice-sensitive DSB task (Wechsler 2013), suggesting practice alone is an unlikely explanation. Another limitation is the low spatial resolution of eLORETA, especially with only 19 EEG channels, which may limit precise source localization despite the strength of within-subject comparisons (Pascual-Marqui 2007). Additionally, behavioral and EEG measures were collected shortly after, but not during, light exposure. Given the dynamic nature of light-induced responses (Vandewalle et al. 2006), future studies should consider real-time assessments and higher-resolution methods such as high-density EEG, MEG, or fMRI.

## Conclusion

A 30-minute session in the light box with exposure to bright, full-spectrum light reduced negative affect, improved cognitive performance, and elicited physiological effects at the EEG level. The most significant changes occurred in the dACC and posterior right insula, regions involved in cognitive control, attention, and emotion regulation. The light intervention had similar behavioral and electrophysiological effects in both SD and ND groups. However, only the SD group exhibited a significant association between increased beta 1 activity and reduced negative affect, indicating a potential mechanism through which light supports emotional regulation. Our findings highlight the potential of bright light interventions to modulate emotional well-being and cognitive function in the adolescent population, which is particularly vulnerable to mood dysregulation and often suffers from partial sleep deprivation. Future research may uncover more targeted applications of light exposure to improve mental health and performance, particularly in individuals with emotional dysregulation.

## Acknowledgements

We sincerely thank the adolescents and their families for their valuable participation and contribution to this study

## Author Contribution

Jana Kopřivová: conceptualization, formal analysis, funding acquisition, methodology, visualization, writing; Zuzana Kaňková: investigation, project administration; Přemysl Vlček: data curation, writing - review & editing; Marek Piorecký: data curation, writing - review & editing; Lenka Maierová: funding acquisition, investigation, writing - review & editing; Zdeňka Bendová: methodology, writing - review & editing; Kateřina Skálová: writing - review & editing; Tereza Nekovářová: conceptualization, methodology, supervision.

## Data Availability Statement

The data that supports the findings of this study are available from the corresponding author, upon reasonable request.

## Generative Artificial Intelligence (AI)

Artificial intelligence-powered tools (ChatGPT version 4.0 and DeepL) were used to improve the language and readability of the manuscript. All AI-generated suggestions were thoroughly reviewed and revised by the authors.

