## Supplementary figures and images for "Bright Light Exposure Reduces Negative Affect and Modulates EEG Activity in Sleep-Deprived and Well-Rested Adolescents"

### Supplementary Figure 1

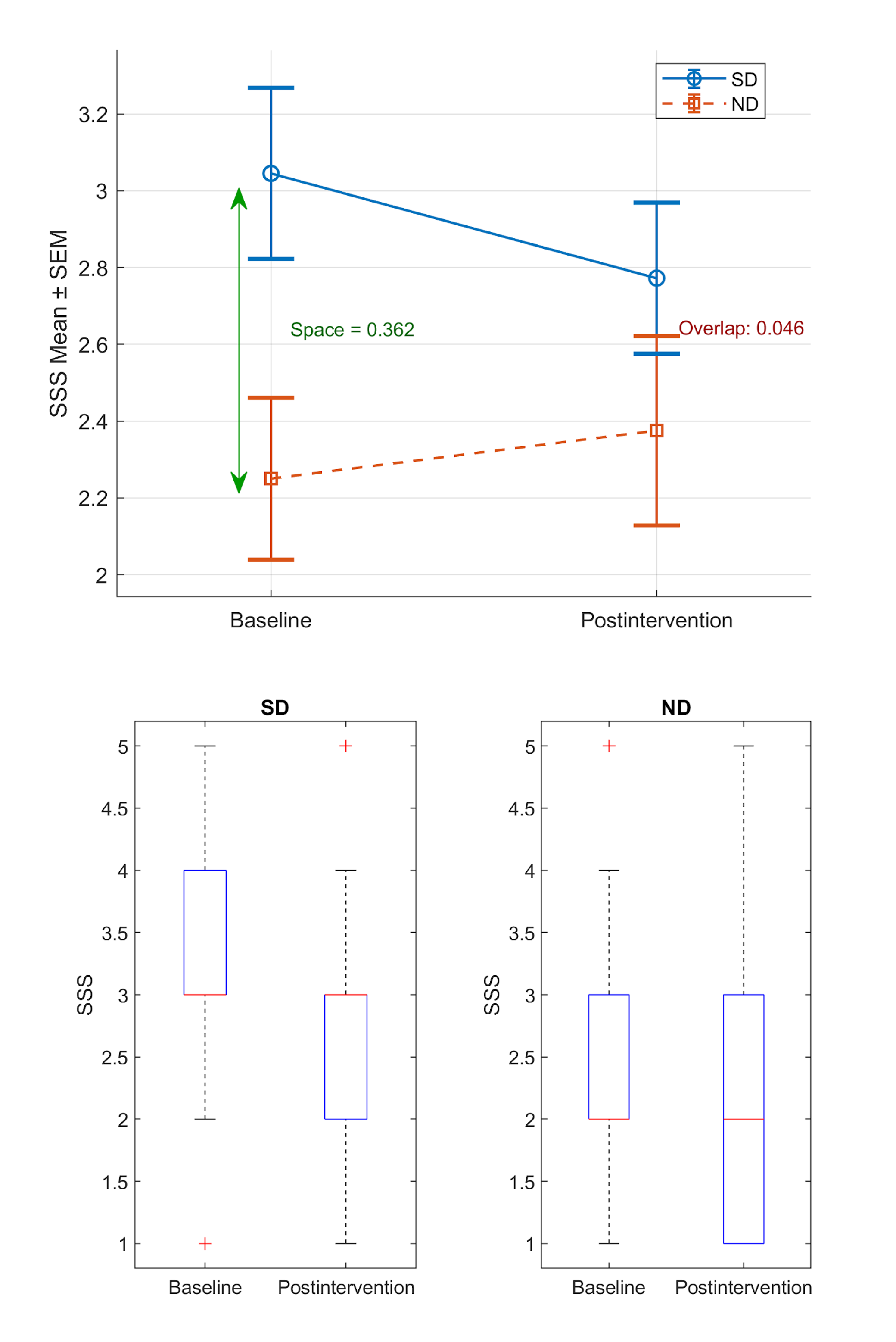
