## Supplementary Figure Legend 1 for "Bright Light Exposure Reduces Negative Affect and Modulates EEG Activity in Sleep-Deprived and Well-Rested Adolescents"

Supplementary Material

**Supplementary Figure 1.** The upper panel shows mean Stanford Sleepiness Scale (SSS) scores at baseline and after the light intervention for the sleep-deprived (SD) and non-sleep-deprived (ND) groups. Error bars represent the standard error of the mean (SEM). Although SSS scores tended to decrease in the SD group following light exposure, this change did not reach statistical significance. The lower panel presents boxplots for each condition, illustrating the distribution and variability of individual SSS scores across participants.
